# Biallelic mutations in *M1AP* are a frequent cause of meiotic arrest leading to male infertility

**DOI:** 10.1101/803346

**Authors:** Margot J. Wyrwoll, Şehime G. Temel, Liina Nagirnaja, Manon S. Oud, Alexandra M. Lopes, Godfried W. van der Heijden, Nadja Rotte, Joachim Wistuba, Marius Wöste, Susanne Ledig, Henrike Krenz, Roos M. Smits, Filipa Carvalho, João Gonçalves, Daniela Fietz, Burcu Türkgenç, Mahmut C. Ergören, Murat Çetinkaya, Murad Başar, Semra Kahraman, Adrian Pilatz, Albrecht Röpke, Martin Dugas, Sabine Kliesch, Nina Neuhaus, GEMINI Consortium, Kenneth I. Aston, Donald F. Conrad, Joris A. Veltman, Corinna Friedrich, Frank Tüttelmann

## Abstract

Male infertility affects ∼7% of men in Western societies, but its causes remain poorly understood. The most clinically severe form of male infertility is non-obstructive azoospermia (NOA), which is, in part, caused by an arrest at meiosis, but so far only few genes have been reported to cause germ cell arrest in males. To address this gap, whole exome sequencing was performed in 60 German men with complete meiotic arrest, and we identified in three unrelated men the same homozygous frameshift variant c.676dup (p.Trp226LeufsTer4) in *M1AP,* encoding meiosis 1 arresting protein. Then, with collaborators from the International Male Infertility Genomics Consortium (IMIGC), we screened a Dutch cohort comprising 99 infertile men and detected the same homozygous variant c.676dup in a man with hypospermatogenesis predominantly displaying meiotic arrest. We also identified two Portuguese men with NOA carrying likely biallelic loss-of-function (LoF) and missense variants in *M1AP* among men screened by the Genetics of Male Infertility Initiative (GEMINI). Moreover, we discovered a homozygous missense variant p.(Pro389Leu) in *M1AP* in a consanguineous Turkish family comprising five infertile men. M1AP is predominantly expressed in human and mouse spermatogonia up to secondary spermatocytes and previous studies have shown that knockout male mice are infertile due to meiotic arrest. Collectively, these findings demonstrate that both LoF and missense *M1AP* variants that impair its protein cause autosomal-recessive meiotic arrest, non-obstructive azoospermia and male infertility. In view of the evidence from several independent groups and populations, *M1AP* should be included in the growing list of validated NOA genes.

## Introduction

Around 7% of all men in Western societies are infertile,^1^ which is primarily diagnosed by semen analysis, comprising as the most relevant parameters sperm concentration and count. More than 10% of all infertile men exhibit azoospermia,^2^ which is defined as the absence of spermatozoa in the ejaculate and constitutes the most clinically severe form of male infertility, resulting in zero chance of natural conception.^3^ Azoospermia is further classified into obstructive and non-obstructive azoospermia (OA and NOA, respectively). In a large fraction of NOA cases a genetic origin is assumed,^4^ such that patients with azoospermia are routinely screened for chromosomal aberrations and Y-chromosomal azoospermia factor (AZF) microdeletions. However, these diagnostic tests only establish a reason for the azoospermia in 15-20% of cases.^2^

NOA can be stratified into four groups based on histological analysis of the seminiferous tubules: normal spermatogenesis (a finding which indicates OA and post-testicular defects), hypospermatogenesis, Sertoli-cell only (SCO), and maturation arrest. Maturation arrest most frequently presents as meiotic arrest, in which spermatocytes are the most advanced germ cell types in the testes.^5^ If germ cell arrest is only partial, some mature spermatozoa will develop, meaning that men with partial germ cell arrest can become parents by undergoing testicular biopsy and sperm extraction (TESE) and then using the extracted sperm in assisted reproductive technology (ART). By contrast, if germ cell arrest is complete, no mature spermatozoa will develop and TESE cannot be successful. However, thus far, germ cell arrest can only be reliably diagnosed by testicular biopsy, i.e., after the surgery, emphasizing the urgent need for better diagnostic tests before biopsy to avoid unnecessary and unsuccessful surgical procedures.

Better diagnostic workup could be developed upon identifying causal genes or mutations that promote germ cell arrest. Recently, the first monogenic alterations associated with germ cell arrest in human males have been described. However, according to a standardized assessment of clinical validity, the X-linked gene *TEX11*^6^ (MIM: 300311) currently remains the only gene with strong evidence.^7^ Given the large number of genes known to cause meiotic arrest in mice, the vast majority of causal mutations causing this phenotype in humans are yet to be identified.

To this end, we first screened the exomes of well-characterized patients with complete, bilateral germ cell arrest at the spermatocyte stage for variants in testis-expressed genes. As a result, we identified biallelic loss-of-function (LoF) variants in three unrelated infertile men in the gene encoding meiosis 1 arresting protein (*M1AP*). Furthermore, by screening two independent cohorts, we found the same homozygous LoF variant in a patient with hypospermatogenesis predominantly displaying meiotic arrest, and we found likely pathogenic missense variants in *M1AP* in two NOA patients. Additionally, we identified a homozygous missense variant in *M1AP* segregating with azoospermia in five infertile men in a consanguineous Turkish family. *M1ap* is primarily expressed in male germ cells during spermatogenesis, and its knockout in male mice leads to infertility due to meiotic arrest.^8, 9^

Our results, together with previously published findings, provide sufficient evidence that M1AP plays an essential role during spermatogenesis and its loss causes NOA in a considerable proportion of infertile men. This allows for a better understanding of the molecular basis of meiotic arrest and improved counseling and treatment of infertile couples.

## Subjects and Methods

### Study cohorts

We originally selected 64 azoospermic but otherwise healthy male patients who attended the Centre of Reproductive Medicine and Andrology (CeRA), University Hospital Münster (N = 51) or the Clinic for Urology, Pediatric Urology and Andrology, Gießen (N = 13), for couple infertility. This is a subset of all patients included in our large-scale Male Reproductive Genomics (MERGE) study, which currently comprises >800 men including 514 with NOA (Figure S1). All of the 64 patients were diagnosed with complete bilateral germ cell arrest at the spermatocyte stage after evaluating at least 100 seminiferous tubules in tissue sections of both testes accompanied by a negative TESE outcome, i.e., no sperm could be recovered.

Chromosomal aberrations and AZF deletions were excluded in advance. Four out of 64 patients were diagnosed with a LoF variant in *TEX11* previously (Yatsenko *et al.* 2015 and unpublished data).^2, 6^ In addition, 27 men, also attending the CeRA for couple infertility and with normal semen parameters (normozoospermia according to WHO^3^), were included as controls. All patients gave written informed consent for the evaluation of their clinical data and analysis of their DNA samples. The study protocol was approved by the respective Ethics Committees/Institutional Review Boards (Ref. No. Münster: 2010-578-f-S, Gießen: 26/11, Nijmegen: NL50495.091.14 version 4, GEMINI consortium: 201502059, Porto: PTDC/SAU-GMG/101229/2008, Bursa: 05.01.2015/04) in accordance with the Helsinki Declaration of 1975.

As a next step, a study cohort of 99 men with unexplained azoospermia (N = 55) or severe oligozoospermia (N = 44) who presented at Radboud University Medical Center (Radboudumc, Nijmegen) was screened for biallelic variants in *M1AP*. In parallel, we screened the whole exome sequencing data produced within the GEMINI study (https://gemini.conradlab.org/) of 979 unrelated men with unexplained NOA for biallelic variants in *M1AP*. In both cohorts, chromosomal aberrations, AZF deletions and *CFTR*-mutations had been excluded.

Additionally, we performed whole exome sequencing in two brothers with unexplained infertility from a consanguineous Turkish family and one fertile brother. The index patient T1024 (V.2; Figure 3A) and his wife presented at Uludag University Faculty of Medicine Hospital because of couple infertility. Semen analysis revealed azoospermia, and chromosomal aberrations as well as AZF deletions were excluded. The patient reported that he had an infertile brother and three further infertile male relatives (Figure 3A). We focused on rare homozygous variants shared by both affected brothers. Subsequently, seven male family members and the mother of T1024 were screened for the *M1AP* variant detected in the infertile brothers.

### Whole exome sequencing (WES) (MERGE study)

Genomic DNA was extracted from peripheral blood leukocytes via standard methods.^10^ WES sample preparation and enrichment were carried out in accordance with the protocols of either Agilent’s SureSelect^QXT^ Target Enrichment kit or Twist Bioscience’s Twist Human Core Exome kit. Agilent’s SureSelect^XT^ Human All Exon Kits V4, V5 and V6 or Twist Bioscience’s Human Core Exome plus RefSeq spike-in’s were used to capture libraries. For multiplexed sequencing, the libraries were index tagged using appropriate pairs of index primers. Quantity and quality of the libraries were assessed with the ThermoFisher Qubit and Agilent’s TapeStation 2200, respectively. Sequencing was conducted on the Illumina HiScan®SQ, NextSeq®500, or HiSeqX® systems using the TruSeq SBS Kit v3 - HS (200 cycles), the NextSeq 500 V2 High-Output Kit (300 cycles) or the HiSeq Rapid SBS Kit V2 (300 cycles), respectively.

After trimming, Cutadapt v1.15 was used to remove the remaining adapter sequences and primers^11^. Sequence reads were aligned against the reference genome GRCh37.p13 using BWA Mem v0.7.17^12^. We excluded duplicate reads and reads that mapped to multiple locations in the genome from further analysis. Small insertions/deletions (indels) and single nucleotide variations were identified and quality-filtered by GATK toolkit v3.8 with HaplotypeCaller, in accordance with the best practice recommendations.^13^ Ensembl Variant Effect Predictor was used to annotate called variants.^14^

DNA extraction, WES and variant calling in patients from the other groups (RU01691, Y126, P86 and T1024) were carried out according to the standard local procedures. The respective details are provided in the Supplemental Methods.

### Data analysis and variant prioritization

In the MERGE study, variation categories, transcript and functional consequences, population frequencies, and *in silico* predicted relevance were annotated to each variant utilizing the in-house pipeline Sciobase^©^. Likely causative variants were identified by filtering the data according to (i) the recessive mode of inheritance, (ii) the population frequency in the Genome Aggregation Database^15^ (gnomAD, minor allele frequency [MAF] < 0.01), and (iii) the functional impact of the variant (loss of function: splice site, frameshift, stop gained/lost, start lost) (Figure S1). Finally, the relevance for the phenotype was assessed using comprehensive expression data (Genotype-Tissue Expression [GTEx] project^16^) and model organism data from the literature. In a complementary approach, an updated version of the population sampling probability (PSAP) pipeline^17^ was used to prioritize potentially causative variants. PSAP models the significance of observing a single subject’s genotype in comparison to genotype frequencies in unaffected populations (commonly referred to as the ‘n-of-one’ problem). The resulting variant lists were filtered as described by Kasak *et al.*^18^ (MAF ≤ 0.01, PopScore ≤ 0.005, and CADD ≥ 20).

To assess the pathogenicity of detected missense variants, we used common *in silico* prediction programs (PolyPhen-2, SIFT, MutationTaster, HOPE^19^). We attempted to model the 3D structure of the M1AP protein. However, due to the lack of previous information on M1AP and comparable 3D structures, it was not possible to achieve a reliable prediction (BLAST for sequence of UniProt identifier Q8TC57 all below 30% sequence identity to known protein structures, details in Suppl. Methods).

### Sanger sequencing for variant validation and screening of normozoospermic controls

All relevant variants identified in azoospermic men were confirmed by direct Sanger sequencing of the respective exons of *M1AP* (NM_001321739.1) according to standard procedures.^10^ To establish the carrier frequency of the recurring variant c.676dup in exon 5 in an ancestry-matched control group, 285 normozoospermic men (from couples attending the CeRA) were analyzed. In the five heterozygous carriers of this variant, the whole coding region of *M1AP* (exons 2-11) was subsequently sequenced, to exclude a second pathogenic variant in *M1AP*. Primer sequences are provided in Table S1.

### Quantitative PCR analysis

To exclude a hemizygous deletion on the other allele, quantitative PCR (qPCR) of exon 5 was performed on gDNA of the three patients from the MERGE study (M330, M864, M1792) carrying the variant c.676dup. qPCR was carried out in 96-well plates on the LightCycler 480 using the manufacturer’s default settings. The *ALB* (albumin) gene was used for normalization. The reactions were performed in triplicates using the SensiMix Real-Time PCR Kit (Bioline). The PCR consisted of an initial incubation step at 95°C for 10 min followed by 40 cycles of 95°C for 15 sec, 60°C for 30 sec and 72°C for 15 sec. Baseline and threshold values were automatically detected using the LightCycler software. Primers are provided in Table S1.

### Histological evaluation and testing of M1AP antibodies

Testis biopsies of patients M330, M864 and M1792 were collected for TESE and research use. Biopsies were fixed in Bouin’s solution overnight at 4°C, washed with 70% ethanol and embedded in paraffin using an automatic ethanol and paraffin row (Bavimed Laborgeräte GmbH, Birkenau, Germany). Subsequently, 5 µm sections were stained with periodic acid-Schiff (PAS) as previously described.^20^

We attempted to establish immunohistochemistry as well as Western blot analyses with commercially available M1AP antibodies (#PA5-31627, ThermoFisher Scientific, Langenselbold, Germany and #HPA045420, Sigma-Aldrich, Munich, Germany). These were selected based on available data from the manufacturers and the human protein atlas (see Suppl. Data for details).

### Search for M1AP variants in women with premature ovarian insufficiency

Because some genes have been reported to be associated with both male and female infertility, we screened 101 women diagnosed with unexplained premature ovarian insufficiency (POI) (62 with isolated POI and 39 with ovarian dysgenesis) by direct Sanger sequencing of the full coding region. Details of a part of this cohort (N = 25) have been published previously.^21^ The additional 76 patients followed the same in- and exclusion criteria.

## Results

### Identification of M1AP as a candidate gene and follow-up study in three independent groups

In the initial MERGE study, the WES data of 60 highly selected, azoospermic infertile patients with unexplained, complete, bilateral germ cell arrest at the spermatocyte stage were screened for rare (MAF < 0.01 according to gnomAD-database^22^), biallelic LoF variants. Two patients were subsequently excluded from this study, because likely causal variants in other genes had been identified in parallel: patient M870 had compound heterozygous variants in *STAG3,*^23^ and patient M1401 had a heterozygous LoF variant in *SYCP2*.^24^ The prioritized genes in the remaining 58 patients were analyzed with regard to the level of expression in the testes. A literature search was performed to identify genes with previous evidence for an association with infertility in either human or model species. The highest prioritized gene was *M1AP* because three unrelated men (M330, M864, M1792, Figure 1) carried the same potentially homozygous LoF variant (c.676dup, MAF = 0.0021, no homozygotes in gnomAD^22^), the M1AP protein displays the highest expression in the testis, and it has been shown to play a crucial role in spermatogenesis in mice.^8, 9^ The variant c.676dup is located in exon 5 of 11 and causes a frameshift and premature stop codon (p.Trp226LeufsTer4), as confirmed by testicular cDNA sequencing of exons 5 of patient M864 (Figure S2B). Quantitative PCR analysis of *M1AP* exon 5 excluded an intragenic deletion and, thus, confirmed homozygosity for c.676dup in all patients (Table S2). No regions of homozygosity (ROH) involving *M1AP* were detected for any of the three patients homozygous for c.676dup, rendering consanguinity of their parents unlikely. Neither did we notice evidence for consanguinity between the patients (analysis by H3M2 algorithm,^25^ data not shown). No patient carrying two rare variants in *M1AP* (MAF ≤ 0.01 from gnomAD) was identified by WES in the remaining MERGE cohort of almost 750 patients with other infertility phenotypes (mostly azoospermia due to other testicular phenotypes such as SCO or without biopsy, N = 548, or severe oligozoospermia, N = 120).

**Figure 1.**
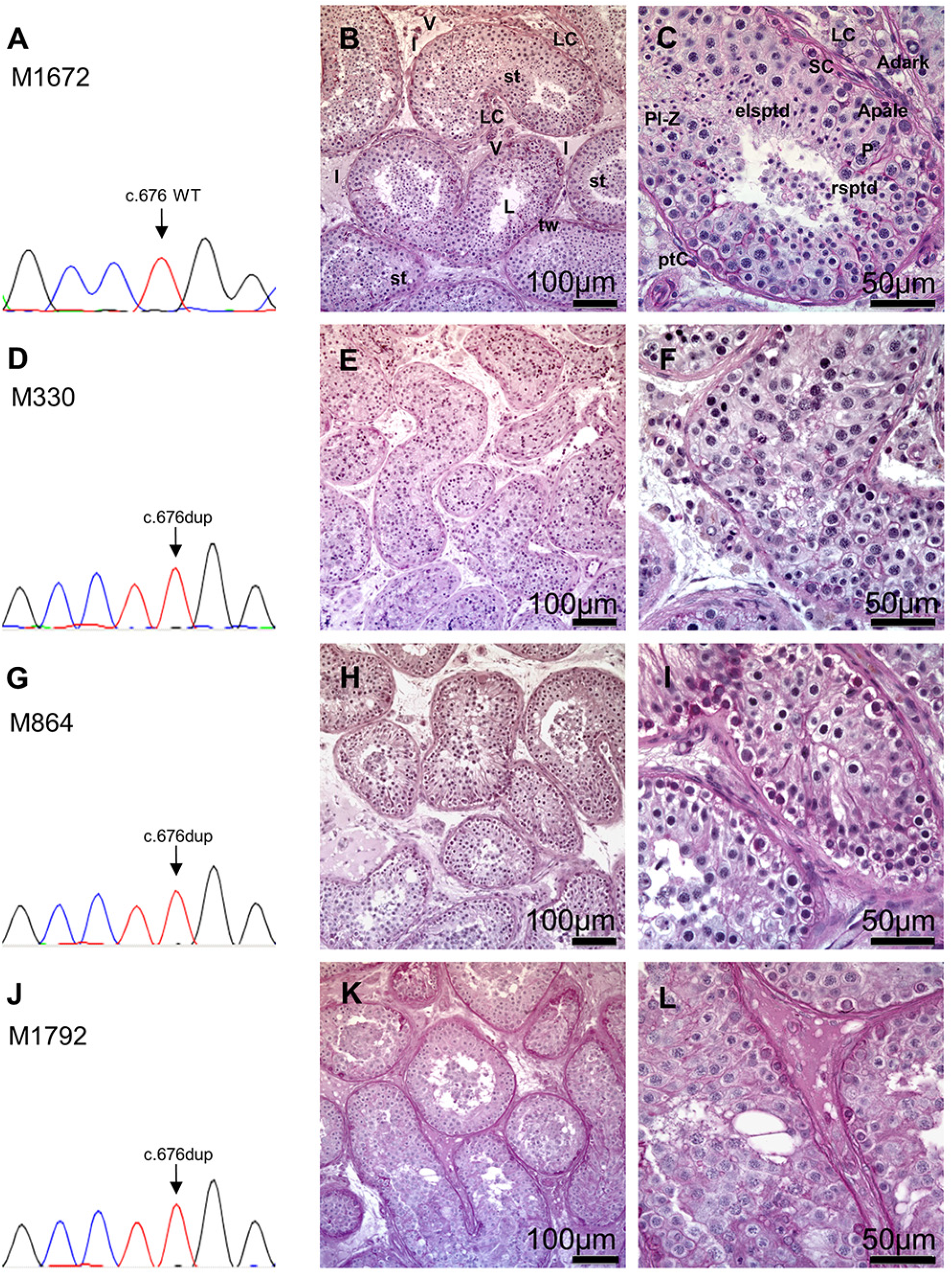
Recurrent homozygous variant c.676dup (duplication of T-nucleotide; p.Trp226LeufsTer4) in *M1AP* leading to complete bilateral meiotic arrest in patients M330 (D-F), M864 (G-I) and M1792 (J-L) from the MERGE study. (**A**) Electropherogram with the wildtype sequence of *M1AP* exon 5 (patient M1672 with obstructive azoospermia). (**B/C**) Testicular tissue showing complete spermatogenesis, PAS staining. (**B**) Testicular tissues are composed of seminiferous tubules and interstitium. The seminiferous tubules are separated from the interstitial space (I) by tubular walls (tw) formed by myoid peritubular cells and the lamina propria. Inside, the seminiferous epithelium and the lumen (L) are localized. In the interstitium, groups of steroidogenic Leydig cells (LC) and blood vessels (V) are observed. Tubular cross section showed the regular appearance of a functioning testis exhibiting complete germ cell differentiation. (**C**) Detail of B; the tubules are surrounded by the lamina propria and the peritubular cells (ptC), forming the wall. Within the seminiferous epithelium, somatic Sertoli cells (SC) are supporting the germ cells differentiating from A spermatogonia (Apale/Adark) via premeiotic spermatocytes (preleptotene to zygotene stage; Pl-Z) into the meiotic pachytene spermatocytes (P). After meiosis is completed, haploid round spermatids (rsptd) are formed which mature further into elongated spermatids (elsptd). (**D-L**) Identification of a recurrent homozygous variant in *M1AP* (c.676dup, p.Trp226LeufsTer4). Sanger sequencing verified the variant in patients M330 (**D**), M864 (**G**) and M1792 (**J**) leading to complete bilateral meiotic arrest as indicated by histological examination of testis biopsies (**E/F**: M330, **H/I**: M864, **K/L**: M1792), which show spermatocytes as most advanced germ cells in all tubules.

Next, and through a collaboration established within the International Male Infertility Genomics Consortium (IMIGC, imigc.org), we identified three additional infertile patients with likely biallelic mutations in *M1AP* from two independent study groups. All identified *M1AP* variants in all six patients were confirmed by Sanger sequencing. The variants, gnomAD frequencies, *in silico* predictions, and clinical data of all patients carrying *M1AP* variants are shown in Table 1.

**Table 1.**
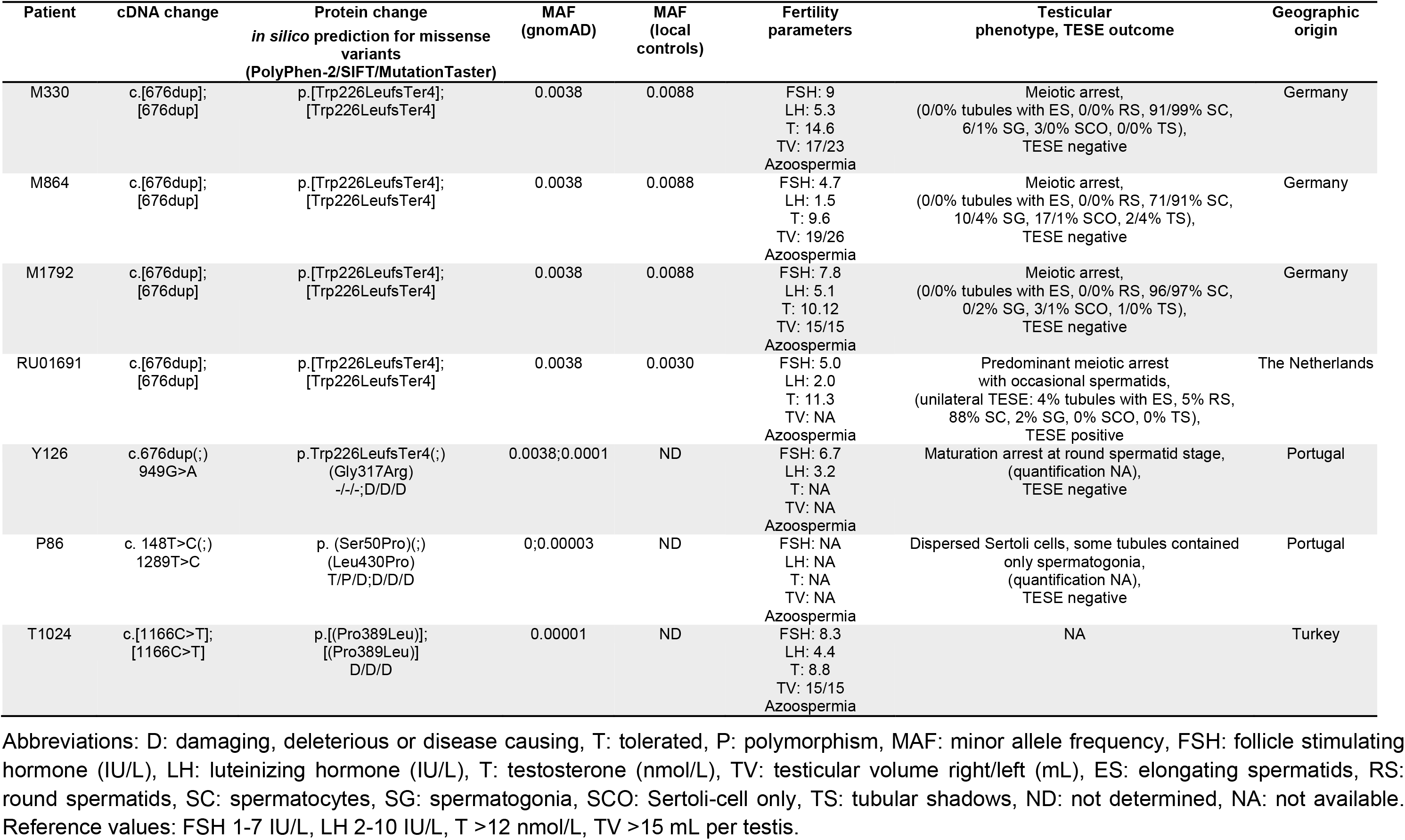
Genetic and clinical data of infertile patients carrying *M1AP* variants.

Patient RU01691 from the Netherlands was also homozygous for the frameshift variant c.676dup (p.Trp226LeufsTer4) in *M1AP* (Figure 2A). The patient’s parents were both heterozygous carriers. Testicular biopsy in this patient showed bilateral severe hypospermatogenesis with predominantly meiotic arrest (Figure 2B); occasionally spermatids were present.

**Figure 2.**
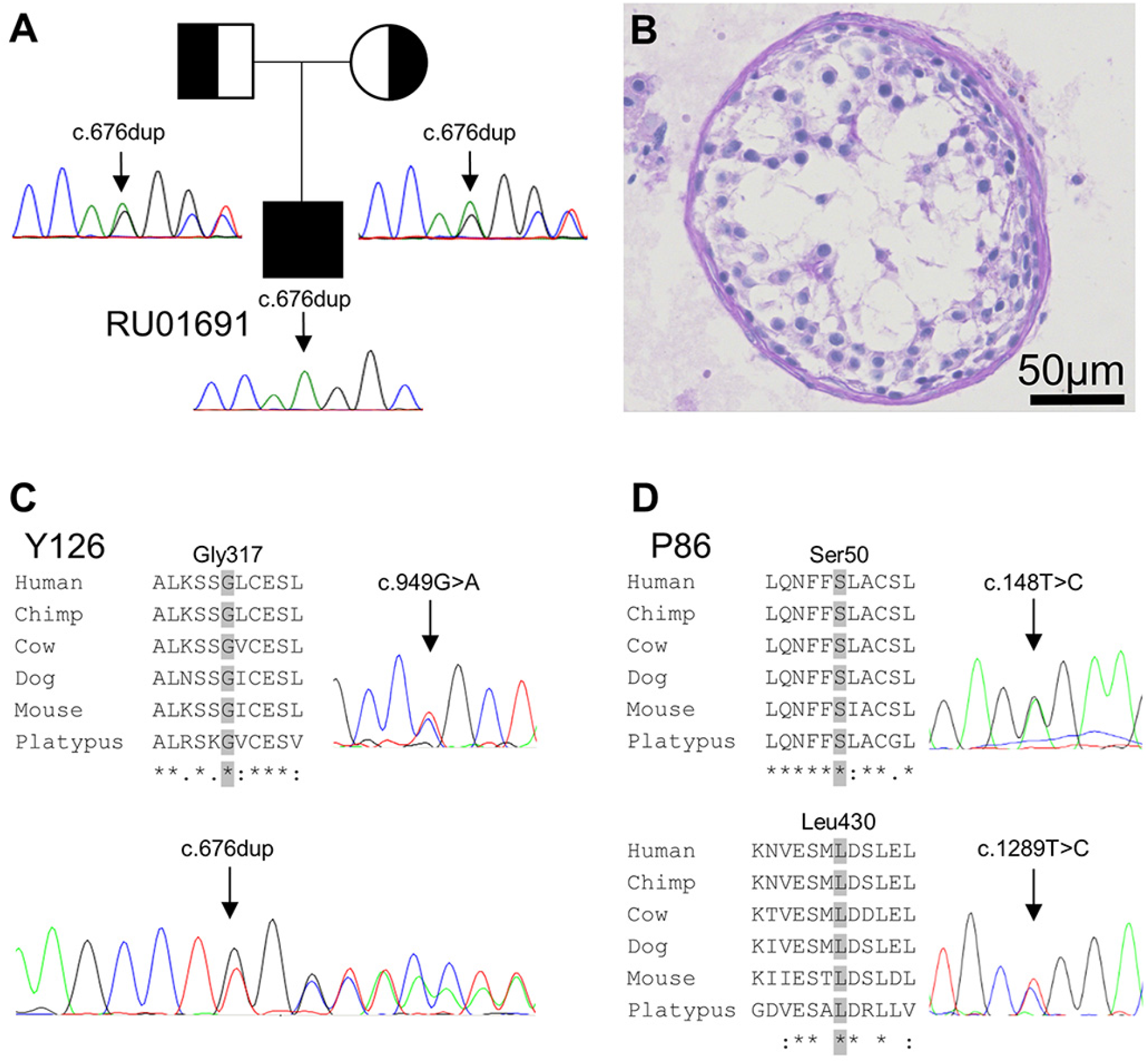
Variants in *M1AP* in two follow-up studies. Recurrent homozygous duplication c.676dup (p.Trp226LeufsTer4) (**A**) in *M1AP* leading to predominantly meiotic arrest in patient RU01691 from Nijmegen, NL. Both parents are heterozygous for the same variant. (**B**) Histology indicates predominant germ cell arrest at the spermatocyte stage. Identification of potentially biallelic variants in *M1AP* in Portuguese patients from the GEMINI study. (**C**) Patient Y126 carries the recurrent LoF variant c.676dup (p.Trp226LeufsTer4) and the missense variant c.949G>A (p.(Gly317Arg)). (**D**) Patient P86 carries two missense variants (c.148T>C(;)1289T>C p.(Ser50Pro)(;)(Leu430Pro)). All missense variants affect highly conserved amino acids, as seen from multiple sequence alignments (**C/D**).

Two patients analyzed within the GEMINI study and of Portuguese origin (P86, Y126) each carried two different variants in *M1AP* (Figure 2C and 2D). Patient Y126 carried the missense variant c.949G>A (p.(Gly317Arg); MAF = 0.00007) and, in addition, the recurrent frameshift variant c.676dup (p.Trp226LeufsTer4). This patient had germ cell arrest at the round spermatid stage as diagnosed by testicular biopsy and histological evaluation. The other patient P86 had two missense variants c.148T>C (p.(Ser50Pro); not described in gnomAD) and c.1289T>C (p.(Leu430Pro); MAF = 0.000008), suggesting compound-heterozygosity. Testicular histology of this patient showed thickened seminiferous tubules and dispersed Sertoli cells with some tubules containing only spermatogonia.

In parallel, and independent of the identification of *M1AP* in the MERGE study, WES was performed in two infertile, azoospermic brothers from a consanguineous Turkish family as well as their fertile brother (Figure 3A). The data were analyzed focusing on rare homozygous variants, which were shared between the infertile brothers but not found in the fertile brother. The two affected men carried rare, homozygous missense variants in the autosomal genes *AMPD2*, *CELSR2*, *CEP164*, and *M1AP* as well as rare hemizygous variants in the X-linked genes *ATG4A* and *ENOX2*. Of these genes, *M1AP* is the only one that has been described in the context of infertility. Both infertile men carried the homozygous missense variant c.1166C>T (p.(Pro389Leu); MAF = 0.00001) in *M1AP*, which was found in a heterozygous state in the fertile brother. No homozygous carriers of this variant have been described so far in any public databases. Subsequently, this variant was also found in a homozygous state in three additional infertile males from the same family. Additionally, after recruiting and examining nine fertile members of the Turkish family, we found that none of them were homozygous for this variant. We did identify both a fertile man and a fertile woman (IV.13 and V.6 in Figure 3A, respectively) as heterozygous carriers of the same variant (example result of Sanger sequencing for subject V.6 shown in Figure 3B). Clinical data was only available for the index case T1024, and this patient had borderline follicle stimulating hormone (FSH, 8.3 U/L) as well as testicular volume (15 mL both left and right) (Table 1). This combination is often found in azoospermic men with germ cell arrest, while most other infertile NOA patients lacking advanced germ cells have elevated FSH levels because of the pituitary-gonadal feedback loop.

**Figure 3.**
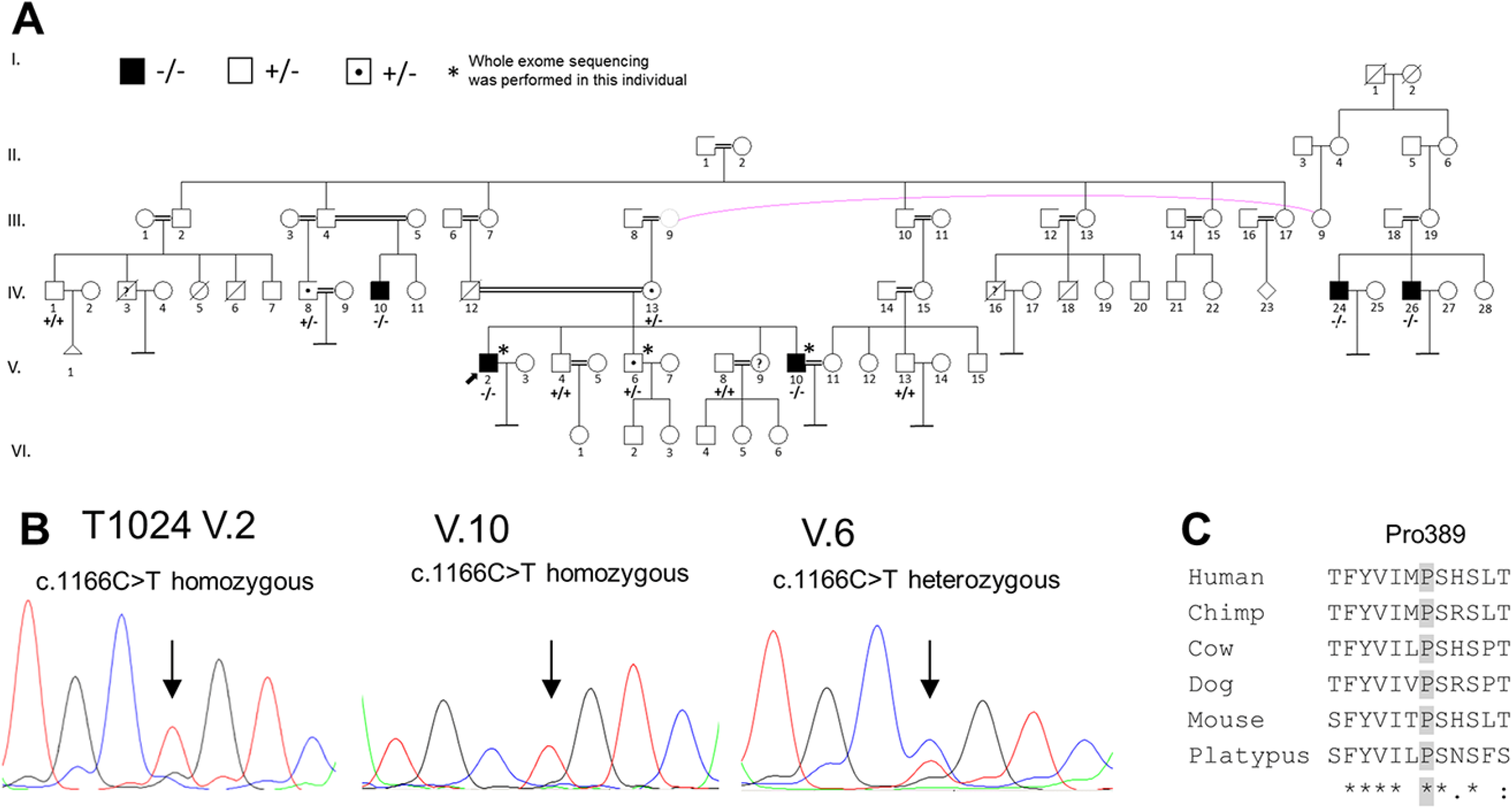
Turkish consanguineous family with infertile, azoospermic men homozygous for *M1AP* missense variant and fertile heterozygous carriers. (**A**) Pedigree of the Turkish family with five infertile azoospermic men carrying the homozygous *M1AP* variant c.1166C>T (p.(Pro389Leu)) indicated with black boxes and -/-. The index patient T1024, who presented at Uludag University Faculty of Medicine Hospital, is marked with an arrow (V.2). Heterozygous carriers of the *M1AP* variant are marked with a point and +/-. Examined family members with the homozygous *M1AP* wildtype allele are marked with +/+. Homozygous carriers are infertile, while heterozygous carriers are fertile. (**B**) Exemplary electropherograms of the index patient (V.2), his infertile brother (V.10), and his fertile brother (V.6), who is a heterozygous carrier. (**C**) The missense variant affects a highly conserved amino acid.

In addition, we analyzed all identified variants from the respective WES analyses in each patient individually utilizing the PSAP pipeline. This pipeline enabled us to rank all variants per patient, following the prioritization criteria (MAF ≤ 0.01, PopScore ≤ 0.005 and CADD ≥ 20). The biallelic *M1AP* LoF variants were ranked in the first position for patient M1792 and in the third position for patients M330 and M864 in the discovery cohort (Table S3). The four patients identified in the follow-up analyses exhibited highly ranked *M1AP* variants as well: position 17 in patient RU01691, position eleven in patient Y126, position 33 in patient P86 and position 12 in T1024, respectively.

### M1AP immunostaining and Western blot

After optimization, both commercial antibodies resulted in a specific signal in immunohistochemistry (Figure S3/4). However, both antibodies seem not to pick up M1AP but a different target. First of all, similar signals could also be detected in two of the patients carrying the homozygous frameshift variant c.676dup, which lead to a disruption of the supposed epitope of the antibodies presumed to reside in the region of or downstream of the variant. Moreover, the Western blot did not only result in detection of a band in testis lysate but also in a kidney lysate where M1AP is not expressed and, last not least, the bands were not at the expected size (Figure S5). Therefore, we also reached out to the colleagues who published the *M1AP* knockout mice and Western blot staining with a self-raised antibody,^9^ but were unsuccessful in getting in contact.

### Control cohorts and women with POI

Because of the rather surprising finding of a rare but recurring variant c.676dup in the primary MERGE study group, we aimed to establish the carrier frequency in an ancestry-matched control population. To this end, we performed Sanger sequencing of exon 5 of *M1AP* in 285 fertile men. We indeed detected five fertile men carrying the same frameshift variant c.676dup (p.Trp226LeufsTer4) in *M1AP*, but these were, in contrast to the three affected patients (M330, M864, M1792), in a heterozygous state. No homozygous carriers were detected, resulting in an allele frequency of 0.0088. We next performed Sanger sequencing for the complete coding region of the *M1AP* gene (exon 2 to 11) in the five heterozygous carriers to rule out the presence of a second variant. No coding alterations were detected.

Additionally, we queried a previously established database of 3347 Dutch fertile couples who had conceived at least one child. WES had been performed in these subjects as part of clinical diagnostics and workup of a child with severe development delay (trio-WES). Again, 20 heterozygous male as well as 10 heterozygous female carriers were detected but no homozygous subjects were found, resulting in an allele frequency of *M1AP* c.676dup in the Dutch cohort of 0.0022. Also, there were no homozygous carriers of other LoF variants among either fathers or mothers. No women carrying two rare variants in *M1AP* were identified in the analyzed 101 women affected by POI.

## Discussion

We have identified a total of eleven patients from four independent cohorts and provide convincing evidence that biallelic variants in *M1AP* are associated with predominantly germ cell arrest in otherwise healthy men. Previously, disruption of *M1ap* has been shown to cause a highly similar testicular phenotype in mice. The *M1ap* knockout mice showed severe oligozoospermia due to predominantly tubular defects with no germ cells beyond the spermatocyte stage, consequently resulting in infertility.^9^ So far, only very few other genes with mutations leading to germ cell arrest in both men and mice and validated in independent cohorts have been published. The first was the X-chromosomal gene *TEX11*,^6, 26^ which is, according to a current structured assessment, one of only a few genes with strong clinical validity for an association with NOA.^7^ Another example is the autosomal gene *STAG3*, which has only recently been described in publications by us and others in parallel,^23, 27^ and its clinical validity is currently ‘moderate’. Most of the proteins involved in DNA recombination, including *M1AP*, are highly evolutionarily conserved.^8^ This suggests that these genes are not tolerant to variation likely due to an infertility phenotype. Because of the previously available evidence from mice and because we replicated our primary finding in several independent groups as well as a consanguineous family, *M1AP* immediately reaches a ‘moderate’ clinical validity (Table S4). This clearly underlines the strength of such collaborative efforts, which have only recently been established in the context of the previously slowly progressing field of male infertility genetics.

From our initial cohort of 64 men with complete bilateral meiotic arrest, we also identified likely causal variants in three other genes, namely *TEX11*, *STAG3* and *SYCP2*. These results have been reported elsewhere.^6, 23, 24^ Among the remaining group of 58 men, we identified three unrelated patients with the same homozygous frameshift variant c.676dup (p.Trp226Leufs*4) in *M1AP*. Thus, up to 5% of men affected by male infertility, non-obstructive azoospermia and meiotic arrest (3 out of 64) carry causal mutations in *M1AP*, making this a comparably common reason for germ cell arrest at the spermatocyte stage like *TEX11* mutations with 6% (4 out of 64) in this selected patient population. Overall, *M1AP* contributes to less than 1% of the highly heterogeneous NOA cases (0.4%, 6 out of 1548 patients in total across the three screened cohorts: 514 and 979 from the MERGE and GEMINI studies, respectively, and 55 from the Nijmegen trio cohort).

The fact that the same frameshift variant c.676dup was also found to be homozygous in RU01691 from the Netherlands and heterozygous in patient Y126 from Portugal suggests that this variant is relatively prevalent in European populations. The likely explanation for its relative commonness is that this mutation is a founder mutation present in men of European ancestry. Yet, although we found this frameshift variant four times in a homozygous state in unrelated men with germ cell arrest, it is a rarely described variant in global large databases. The gnomAD database does not list any homozygous men, and the maximum allele frequency is 0.0038 in the European (non-Finnish) population. In our control cohort of 285 German men with normal sperm production, we detected five heterozygous carriers of c.676dup, corresponding to an allele frequency of 0.0088. This difference in allele frequency can be explained either by the relatively small size of the control cohort or by an enrichment in the population attending the CeRA in Münster, i.e., of Westphalian origin. In the larger Dutch cohort of 3347 fertile couples, 30 heterozygous carriers were found, corresponding to an allele frequency of 0.0022.

The relevance of the homozygous frameshift variant c.676dup in *M1AP* found in four patients is underlined by the very low PopScore (9.7×10^-7^) obtained by PSAP and the high prioritization of the *M1AP* variants in all analyzed patients who carried the variants (Table S3). Moreover, the expected mode of inheritance for this gene is autosomal recessive according to a general prediction,^28^ fitting our observations of biallelic variants in the affected patients.

This frameshift variant c.676dup very likely causes a premature stop codon four amino acids downstream (p.Trp226Leufs*4). This could result either in an altered and severely truncated protein with 230 amino acids (normal protein length: 530 amino acids), or in nonsense-mediated decay (NMD) of the mRNA. Analysis of patient M864’s testis RNA resulted in an equal band compared to control testis RNA, and we therefore exclude elimination through NMD (Figure S2A). Unfortunately, both protein analysis via Western blot and immunohistological stainings failed because of the lack of suitable antibodies raised against M1AP (Figures S3, S4, and S5).

In addition to the heterozygous frameshift c.676dup, patient Y126 carries the substitution c.949G>A. The missense variant replaces the highly conserved (up to platypus; Figure 2C) neutral and nonpolar amino acid glycine with the larger, positively charged amino acid arginine (p.(Gly317Arg)). Based on conservation information, the variation in this position is highly likely to impair M1AP protein function.^19^ The introduction of a charge may cause the repulsion of interaction partners or of other positively charged residues. In addition, the altered torsion angles may have an influence on the correct conformation and disturb the local structure of the protein. Correspondingly, the amino acid change p.(Gly317Arg) is predicted to be deleterious by all *in silico* algorithms.

Patient P86 carries two missense variants in *M1AP*. The mutation c.1289T>C leads to a substitution of the hydrophobic amino acid leucine, which is predicted to be located in an alpha helix, with the less hydrophobic amino acid proline (p.(Leu430Pro)).^19^ Because proline is an alpha helix breaker, the alteration likely has severe effects on protein structure. This change is predicted to be disease causing by all *in silico* programs. Furthermore, leucine at position 430 is a highly conserved amino acid (up to platypus; Figure 2D). The substitution c.148T>C replaces the polar amino acid serine at position 50 with the nonpolar amino acid proline (p.(Ser50Pro)). Again, the wildtype residue is predicted to be located in an alpha helix, and, therefore, its substitution by a proline likely has severe effects on protein structure and function.^19^ Although the *in silico* algorithms SIFT and MutationTaster predict this change as being tolerated and as a polymorphism, the variant has not previously been described in any population, which supports a pathogenic impact. Moreover, this amino acid is likewise highly conserved (up to platypus; Figure 2D).

We add another layer of evidence that *M1AP* missense variants cause NOA with data from a consanguineous family showing segregation of the homozygous *M1AP* missense variant c.1166C>T (p.(Pro380Leu) with affected family members (LOD score = 3.28). Of note, the group of SGM identified *M1AP* as candidate for the affected, azoospermic family members independently from our initial identification. Although the index case T1024 did not undergo a testicular biopsy, his clinical data is indicative of NOA due to germ cell arrest. In contrast, all fertile family members investigated had at least one wildtype allele, which underlines the pathogenicity of this variant when it is biallelic. The substitution c.1166C>T leads to a replacement of the highly conserved and weakly hydrophobic proline with the more hydrophobic leucine at position 389 (p.(Pro389Leu), Figure 3C). Proline is known to have a rigid structure, giving a protein a specific conformation, which could be disrupted by substitution with leucine.^19^ Accordingly, the amino acid exchange is predicted to be pathogenic by all *in silico* programs.

The possible effects of the detected missense variant on post-translational modifications and folding of the M1AP protein are difficult to assess and, thus, inherently uncertain. Unfortunately, the structure of the M1AP protein is currently unknown and it is impossible to predict functional domains of the M1AP protein with high reliability. Moreover, the specific molecular function of M1AP remains to be elucidated in subsequent studies.

The process of meiosis is in part similar between males and females, but orchestrated highly differently concerning its timing. Thus far, only few genes have been reported to impair male as well as female meiosis. As an example, variants in *STAG3* had previously been reported to cause POI^29^ and now have recently been shown to also cause NOA and meiotic arrest.^23, 27^ *M1AP* is predominantly expressed in the adult testis (GTEx), but also reported to be expressed in the fetal mouse ovary.^8^ However, the structure of ovaries in female *M1ap* knockout mice appeared normal and fertility was preserved.^9^ Concordantly, we did not identify any relevant variants in *M1AP* in 101 women affected by POI. Still, we neither found a fertile woman carrying a homozygous LoF variant in the large Dutch trio cohort. From this data, we cannot exclude that variants in *M1AP* may be a rare cause also for POI, but it seems likely that M1AP is only required for male meiosis.

The common phenotype among our patients is NOA, and four out of six with testicular histology available had either complete (N = 3) or predominant (N = 1) meiotic arrest, i.e., germ cell arrest at the spermatocyte stage. A similar phenotype was observed in mice with disruption of *M1ap*,^9^ but these mice had some sperm in their semen, i.e., severe oligozoospermia. This fact is not inconsistent with our findings, as the authors of the study described mice exhibiting variable efficiency of *M1ap* disruption: Some mice had no apparent wildtype protein, while others displayed reduced levels of the M1AP protein.^9^ If the authors had achieved a complete biallelic knockout of *M1ap* in all mice, representing complete meiotic arrest, one would expect azoospermia to result.

We did not identify biallelic *M1AP* variants in any other male infertility phenotypes such as SCO or severe oligozoospermia, and no subjects with homozygous *M1AP* LoF variants are present in gnomAD (>140,000 subjects). In conclusion, the presented data strongly suggests that both homozygous LoF and missense variants in *M1AP* impairing its protein as well as compound heterozygosity for either variant type lead to NOA. *M1AP* disruption is associated primarily with germ cell arrest at meiosis/spermatocyte stage, but it may also be compatible with some rare instances of further progressed spermatogenesis. Based on our data, we cannot reliably predict the probability of successful sperm retrieval by testicular biopsy and TESE among men with disrupted *M1AP*, but if we extrapolate the findings from the men reported here, TESE success is quite low (1 of 6, <20%).

Our finding that biallelic *M1AP* mutations cause predominantly germ cell arrest at the spermatocyte stage in infertile men provides further evidence that meiotic arrest is often of monogenic origin. According to the structured assessment presented here, *M1AP* has considerable clinical validity for causing NOA and could potentially be used as a screening marker before testicular biopsy to estimate the chances of successful TESE. Last not least, identifying mutations in *M1AP* in infertile NOA men provides them with a causal diagnosis for their infertility.

## Appendices

### Supplemental Data

Supplemental Data includes five tables, five figures, and supplemental methods.

## Acknowledgements

We are indebted to all patients consenting to research evaluation of their data and donating their DNA as well as the physicians who took care of them. We thank Christian Ruckert for his bioinformatic support, Martin Bergmann for many years of skillfully evaluating testicular histologies, Zeliha Görmez for bioinformatics analyses of WES, Şeref Gül and Laurens van de Wiel for their attempts to model the M1AP protein, Joachim Kremerskothen and Verena Höffken for attempting to establish a Western blot analysis, as well as Christina Burhöi, Nicole Terwort, and Katja Hagen for their excellent technical assistance. We thank Dr. Celeste Brennecka for language editing of the manuscript.

This work was carried out within the frame of the German Research Foundation Clinical Research Unit ‘Male Germ Cells: from Genes to Function’ (DFG CRU326). Funding for sequencing of the GEMINI cohort was provided by the National Institutes of Health (R01HD078641).The analyses in the Turkish family were supported by a grant from the Bursa University of Uludag Project Unit (KUAP(T)-2014/36).

## Web Resources

Clinical Research Unit ‘Male Germ Cells’, http://www.male-germ-cells.de

ClinVar, https://www.ncbi.nlm.nih.gov/clinvar

GEMINI, https://gemini.conradlab.org/

gnomAD, https://gnomad.broadinstitute.org

GTEx, https://gtexportal.org

HOPE, https://www3.cmbi.umcn.nl/hope/input/

Human Protein Atlas, https://www.proteinatlas.org/

Male Fertility Gene Atlas, https://mfga.uni-muenster.de

MutationTaster, http://www.mutationtaster.org/

International Male Infertility Genomics Consortium, http://www.imigc.org

OMIM, http://www.omim.org

PolyPhen 2, http://genetics.bwh.harvard.edu/pph2/

PSAP, https://github.com/conradlab/PSAP

SIFT, https://sift.bii.a-star.edu.sg/

## Accession Numbers

All variants have been submitted to ClinVar (SUB6396814 [*will be substituted for final accession no. when available*]) and can also be accessed in the Male Fertility Gene Atlas (MFGA, https://mfga.uni-muenster.de), a public platform for collecting and integrating data sets about epi-/genetic causes of male infertility produced in a subproject of the Clinical Research Unit ‘Male Germ Cells: from Genes to Function’.

## References

1. WHO | World Health Organization (2017). Sexual and reproductive health - Fertility and infertility - Assisting couples and individuals.

2. Tüttelmann, F., Ruckert, C., and Röpke, A. (2018). Disorders of spermatogenesis: Perspectives for novel genetic diagnostics after 20 years of unchanged routine. Medizinische Genet. 30, 12–20.

3. WHO | World Health Organization (2010) WHO laboratory manual for the examination and processing of human semen.

4. Lee, J.Y., Dada, R., Sabanegh, E., Carpi, A., and Agarwal, A. (2011). Role of Genetics in Azoospermia. URL 77, 598–601.

5. Matzuk, M.M., and Lamb, D.J. (2008). The biology of infertility: Research advances and clinical challenges. Nat. Med. 14, 1197–1213.

6. Yatsenko, A.N., Georgiadis, A.P., Röpke, A., Berman, A.J., Jaffe, T., Olszewska, M., Westernströer, B., Sanfilippo, J., Kurpisz, M., Rajkovic, A., et al. (2015). X-Linked *TEX11* Mutations, Meiotic Arrest, and Azoospermia in Infertile Men. N. Engl. J. Med. 372, 2097– 2107.

7. Oud, M.S., Volozonoka, L., Smits, R.M., Vissers, L.E.L.M., Ramos, L., and Veltman, J.A. (2019). A systematic review and standardized clinical validity assessment of male infertility genes. Hum. Reprod. 34, 932–941.

8. Arango, N.A., Huang, T.T., Fujino, A., Pieretti-Vanmarcke, R., and Donahoe, P.K. (2006). Expression analysis and evolutionary conservation of the mouse germ cell-specific D6Mm5e gene. Dev. Dyn. 235, 2613–2619.

9. Arango, N.A., Li, L., Dabir, D., Nicolau, F., Pieretti-Vanmarcke, R., Koehler, C., McCarrey, J.R., Lu, N., and Donahoe, P.K. (2013). Meiosis I Arrest Abnormalities Lead to Severe Oligozoospermia in Meiosis 1 Arresting Protein (M1ap)-Deficient Mice1. Biol. Reprod. 88, 1– 11.

10. Röpke, A., Tewes, A.-C., Gromoll, J., Kliesch, S., Wieacker, P., and Tüttelmann, F. (2013). Comprehensive sequence analysis of the NR5A1 gene encoding steroidogenic factor 1 in a large group of infertile males. Eur. J. Hum. Genet. 21, 1012–1015.

11. Martin, M. (2014). Cutadapt removes adapter sequences from high-throughput sequencing reads. EMBnet.journal 17, 10.

12. Li, H., and Durbin, R. (2010). Fast and accurate long-read alignment with Burrows – Wheeler transform. Bioinformatics 26, 589–595.

13. McKenna, A., Hanna, M., Banks, E., Sivachenko, A., Cibulskis, K., Kernytsky, A., Garimella, K., Altshuler, D., Gabriel, S., Daly, M., et al. (2010). The genome analysis toolkit: A MapReduce framework for analyzing next-generation DNA sequencing data. Genome Res. 20, 1297–1303.

14. McLaren, W., Gil, L., Hunt, S.E., Riat, H.S., Ritchie, G.R.S., Thormann, A., Flicek, P., and Cunningham, F. (2016). The Ensembl Variant Effect Predictor. Genome Biol. 17,.

15. Karczewski, K.J., Francioli, L.C., Tiao, G., Cummings, B.B., Alföldi, J., Wang, Q., Collins, R.L., Laricchia, K.M., Ganna, A., Birnbaum, D.P., et al. (2019). Variation across 141,456 human exomes and genomes reveals the spectrum of loss-of-function intolerance across human protein-coding genes. bioRxiv 531210.

16. Lonsdale, J., Thomas, J., Salvatore, M., Phillips, R., Lo, E., Shad, S., Hasz, R., Walters, G., Garcia, F., Young, N., et al. (2013). The Genotype-Tissue Expression (GTEx) project. Nat. Genet. 45, 580.

17. Wilfert, A.B., Chao, K.R., Kaushal, M., Jain, S., Zöllner, S., Adams, D.R., and Conrad, D.F. (2016). Genome-wide significance testing of variation from single case exomes. Nat. Genet. 48, 1455–1461.

18. Kasak, L., Punab, M., Nagirnaja, L., Grigorova, M., Minajeva, A., Lopes, A.M., Punab, A.M., Aston, K.I., Carvalho, F., Laasik, E., et al. (2018). Bi-allelic Recessive Loss-of-Function Variants in FANCM Cause Non-obstructive Azoospermia. Am. J. Hum. Genet. 103, 200–212.

19. Venselaar, H., te Beek, T.A., Kuipers, R.K., Hekkelman, M.L., and Vriend, G. (2010). Protein structure analysis of mutations causing inheritable diseases. An e-Science approach with life scientist friendly interfaces. BMC Bioinformatics 11, 548.

20. Nieschlag, E., Behre, H.M., and Nieschlag, S. (2010). Andrology: Male reproductive health and dysfunction.

21. Ledig, S., Röpke, A., and Wieacker, P. (2010). Copy number variants in premature ovarian failure and ovarian dysgenesis. Sex. Dev. 4, 225–232.

22. Lek, M., Karczewski, K.J., Minikel, E. V, Samocha, K.E., Banks, E., Fennell, T., O’Donnell-Luria, A.H., Ware, J.S., Hill, A.J., Cummings, B.B., et al. (2016). Analysis of protein-coding genetic variation in 60,706 humans. Nature 536, 285.

23. van der Bijl, N., Röpke, A., Biswas, U., Wöste, M., Jessberger, R., Kliesch, S., Friedrich, C., and Tüttelmann, F. (2019). Mutations in STAG3 cause male infertility due to meiotic arrest. Hum. Reprod.

24. Schilit, S.L.P., Menon, S., Friedrich, C., Kammin, T., Wilch, E., Hanscom, C., Jiang, S., Kliesch, S., Talkowski, M.E., Tüttelmann, F., et al. (2019). SYCP2 translocation-mediated dysregulation and frameshift variants cause human male infertility. bioRxiv 641928.

25. Magi, A., Tattini, L., Palombo, F., Benelli, M., Gialluisi, A., Giusti, B., Abbate, R., Seri, M., Gensini, G.F. ranc., Romeo, G., et al. (2014). H3M2: detection of runs of homozygosity from whole-exome sequencing data. Bioinformatics 30, 2852–2859.

26. Yang, F., Silber, S., Leu, N.A., Oates, R.D., Marszalek, J.D., Skaletsky, H., Brown, L.G., Rozen, S., Page, D.C., and Wang, P.J. (2015). TEX11 is mutated in infertile men with azoospermia and regulates genome-wide recombination rates in mouse. EMBO Mol. Med. 7, 1198–1210.

27. Riera-Escamilla, A., Enguita-Marruedo, A., Moreno-Mendoza, D., Chianese, C., Sleddens-Linkels, E., Contini, E., Benelli, M., Natali, A., Colpi, G.M., Ruiz-Castañé, E., et al. (2019). Sequencing of a “mouse azoospermia” gene panel in azoospermic men: identification of RNF212 and STAG3 mutations as novel genetic causes of meiotic arrest. Hum. Reprod. 34, 978–988.

28. Cassa, C.A., Weghorn, D., Balick, D.J., Jordan, D.M., Nusinow, D., Samocha, K.E., O’Donnell-Luria, A., MacArthur, D.G., Daly, M.J., Beier, D.R., et al. (2017). Estimating the selective effects of heterozygous protein-truncating variants from human exome data. Nat. Genet. 49, 806–810.

29. Caburet, S., and Vilain, É. (2015). STAG3 in premature ovarian failure. Medecine/Sciences 31, 129–131.

