## Supplemental Material for "Biallelic mutations in *M1AP* are a frequent cause of meiotic arrest leading to male infertility"

**Table S1. Primer sequences.**

| Primer | Sequence Forward | Sequence Reverse |
|---|---|---|
| Validation of c.676dup (MERGE study) | TGGGTCTGGAAATGTTGCTGA | GATTGCTAGAGCCCAGGCAT |
| qPCR of Exon 5 in *M1AP* | TCTGGGAACTGACATTGACCTTC | TGGGTCTGGAAATGTTGCTGA |
| Sanger-Sequencing of all exons of *M1AP* in heterozygous control men | | |
| Exon 2 | TGGATTTTCTCTTCAACAGTACACA | TGTGCACCTGTAGTCCTAGC |
| Exon 3 | CAGTTTTTCCTCATAATTCACTTCAGT | TCCTTTCATGTTTCTGGTAACTCT |
| Exon 4 | CCTCAGTGAAATCTGCTGGC | GCCTGATTGGAAAGGTCCTGT |
| Exon 5 | CTAACTGGCCCTTGCTGGTT | TGCTAGAGCCCAGGCATTTG |
| Exon 6 | CACCATCTGCACATTTGGCC | AACCAGTCAGGCTTTCCTCT |
| Exon 7 | ACAGAATATATATCTAGGGCTTGACAC | GAGTCTGCTTCAACTCTTCCCA |
| Exon 8 | GCCGAAGTTAAATGGCTCTG | TGGCATAATTGCCTATCCTT |
| Exon 9+10 | GGGGGACAGCATCTATTTCA | TTCCCTCTTCAACCCCAACT |
| Exon 11 | CCTTGAGGCTGTCACTCCA | CTTGCTGGAGAAAGGACAGG |
| RNA analyses | | |
| M1AP cDNA Ex2-3 | ACATTGCTCTACCGTCCTGG | TGCAACCTAGCAAAGTTCCCT |
| M1AP cDNA Ex4-6 | CCTAGCCAGAGTCAGGAGGT | ATTCTCAAGGAGCCGTCAGC |

**Table S2. qPCR results to exclude the hemizygosity of LoF variant c.676dup in exon 5 of *M1AP* of patients from the MERGE study.**

| Patient | Repeat | Crossing Point Exon 5 *M1AP* | Concentration Exon 5 *M1AP* | Mean concentration Exon 5 *M1AP* | Crossing Point *Albumin* | Concentration *Albumin* | Mean concentration *Albumin* | Ratio mean concentration Exon 5 *M1AP* / mean concentration *Albumin* |
|---|---|---|---|---|---|---|---|---|
| M330 | 1 | 21.58 | 10.2 | | 22.64 | 9.42 | | |
| M330 | 2 | 21.57 | 10.3 | 10.03 | 22.65 | 9.35 | 9.40 | 1.10 |
| M330 | 3 | 21.57 | 10.3 | | 22.64 | 9.43 | | |
| M864 | 1 | 22.62 | 4.96 | | 23.46 | 5.34 | | |
| M864 | 2 | 22.58 | 5.12 | 5.04 | 23.46 | 5.34 | 5.35 | 0.94 |
| M864 | 3 | 22.60 | 5.03 | | 23.45 | 5.37 | | |
| M1792 | 1 | 21.96 | 7.86 | | 23.21 | 6.37 | | |
| M1792 | 2 | 21.97 | 7.79 | 7.80 | 23.12 | 6.77 | 6.78 | 1.16 |
| M1792 | 3 | 21.98 | 7.73 | | 23.03 | 7.20 | | |
| Control | 1 | 21.62 | 9.95 | | 22.54 | 10.0 | | |
| Control | 2 | 21.61 | 10.0 | 10.0 | 22.55 | 10.0 | 10.0 | 1.0 |
| Control | 3 | 21.60 | 10.0 | | 22.57 | 10.0 | | |

**Table S3. Top 50-List of PSAP results of patients with *M1AP* variants.**

(see respective Excel-file)

**Table S4. Structured clinical validity assessment of M1AP according to Smith et al. 2018.**

| | Points | Comments/Reference |
|---|---|---|
| **Number of unrelated patients** (1-2 → 1 pt, 3-4 → 2 pt, 5-9 → 3 pt, 10-24 → 4 pt; > 25 → classification "definitive") | 3 | 7 patients (M330, M864, M1792, RU01691, Y126, P86, family of T1024) |
| **Other statistical evidence** (AD disease with significant excess of de novos OR AR disease with e.g. LOD score > 3 → 1 pt) | 1 | consanguineous Turkish family (homozygous variant segregates with infertility, heterozygous carriers fertile, LOD score 3.28) |
| **Number of publications reporting independent probands** (per publication 1 pt, max. 3) | 1 (2) | this publication/manuscript; in brackets: if we had published the findings in the Nijmegen patient separately |
| **Number of pathogenic variants** (per VLP or mutation 1 pt, max. 4) | 1 | c.676dup (p.Trp226LeufsTer4) |
| **Gene function** (function/expression consistent with disease → 1 pt | 1 | Arango et al. 2006: testicular expression in mice, expressed in last stages of spermatogenesis, Human Protein Atlas, GTEx |
| …and/or physically interacts with gene characterized for same disease → 1 pt) | 0 | |
| **Gene disruption** (relevant pathology in vitro after similar genetic modification → 1 pt | 0 | |
| ...and/or determination of mutational mechanism → 1 pt) | 1 | Frameshift variant → truncated protein → LoF (this study: analysis of testis RNA) |
| **Model organism** (gene function in vivo related to pathology of human disease → 1 pt | 1 | Arango et al. 2013: male KO-mouse is infertile |
| …and/or phenotype and genotype match human disease → 1 pt) | 1 | Arango et al. 2013: male knockout mouse exhibits meiotic arrest |
| **TOTAL POINTS** | 10 (11) | |
| **Classification**<br>no evidence (0-4 pts)<br>limited (2-9 pts)<br>moderate (8-12 pts)<br>strong (13+ pts)<br>definitive (canonical) | moderate | |

**Table S5. Members of the GEMINI consortium.**

| **Group Leaders** | **Institution** |
|---|---|
| Donald F. Conrad, Liina Nagirnaja | Department of Genetics, Oregon National Primate Research Center, Oregon Health & Science University, Beaverton, OR, USA |
| Kenneth I. Aston, Douglas T. Carrell, James M. Hotaling, Timothy G. Jenkins | Andrology and IVF Laboratory, Department of Surgery (Urology), University of Utah School of Medicine, Salt Lake City, UT, USA |
| Rob McLachlan[1,2] | 1) Hudson Institute of Medical Research and the Department of Obstetrics and Gynaecology, Monash University, Clayton, Victoria, Australia<br>2) Monash IVF and the Hudson Institute of Medical Research, Clayton, Victoria, Australia |
| Moira K. O'Bryan | School of Biological Sciences, Monash University, Clayton, Victoria, Australia |
| Peter N. Schlegel | Department of Urology, Weill Cornell Medicine, New York, NY, USA |
| Michael L. Eisenberg | Department of Urology, Stanford University School of Medicine, Stanford, CA 94305, USA |
| Jay I. Sandlow | Department of Urology, Medical College of Wisconsin, Milwaukee, WI, 53226, USA |
| Emily S. Jungheim, Kenan R. Omurtag | Washington University in St Louis, School of Medicine, St Louis, MO, USA |
| Alexandra M. Lopes[1,2], Susana Seixas[1,2], Filipa Carvalho[1,3], Susana Fernandes[1,3], Alberto Barros[1,3] | 1) i3S - Instituto de Investigação e Inovação em Saúde , Universidade do University of Porto<br>2) IPATIMUP - Instituto de Patologia e Imunologia Molecular da Universidade do Porto, Porto, Portugal<br>3) Serviço de Genética, Departamento de Patologia, Faculdade de Medicina da Universidade do Porto, Porto, Portugal |
| João Gonçalves[1,2], Iris Caetano[1], Graça Pinto[3], Sónia Correia[3] | 1) Departamento de Genética Humana, Instituto Nacional de Saúde Dr Ricardo Jorge, Lisboa, Portugal<br>2) ToxOmics, Faculdade de Ciências Médicas, Universidade Nova de Lisboa, Portugal<br>3) Centro de Medicina Reprodutiva, Maternidade Dr. Alfredo da Costa, Lisboa, Portugal |
| Maris Laan | Institute of Biomedicine and Translational Medicine, University of Tartu, 51010 Tartu, Estonia |
| Margus Punab | Andrology Center, Tartu University Hospital, 50406 Tartu, Estonia |
| Ewa Rajpert-De Meyts, Niels Jørgensen, Kristian Almstrup | Department of Growth and Reproduction, Rigshospitalet, University of Copenhagen, Copenhagen, Denmark |
| Csilla G. Krausz[1,2] | 1) Department of Experimental and Clinical Biomedical Sciences, University of Florence, Florence, Italy<br>2) Andrology Department, Fundacio Puigvert, Instituto de Investigaciones Biomédicas Sant Pau (IIB-Sant Pau), Barcelona, Spain |
| Keith A. Jarvi | Division of Urology, Department of Surgery, Mount Sinai Hospital and; Institute of Medical Sciences, University of Toronto, Toronto, Ontario, Canada |

**Figure S1. Prioritization scheme for WES data.**

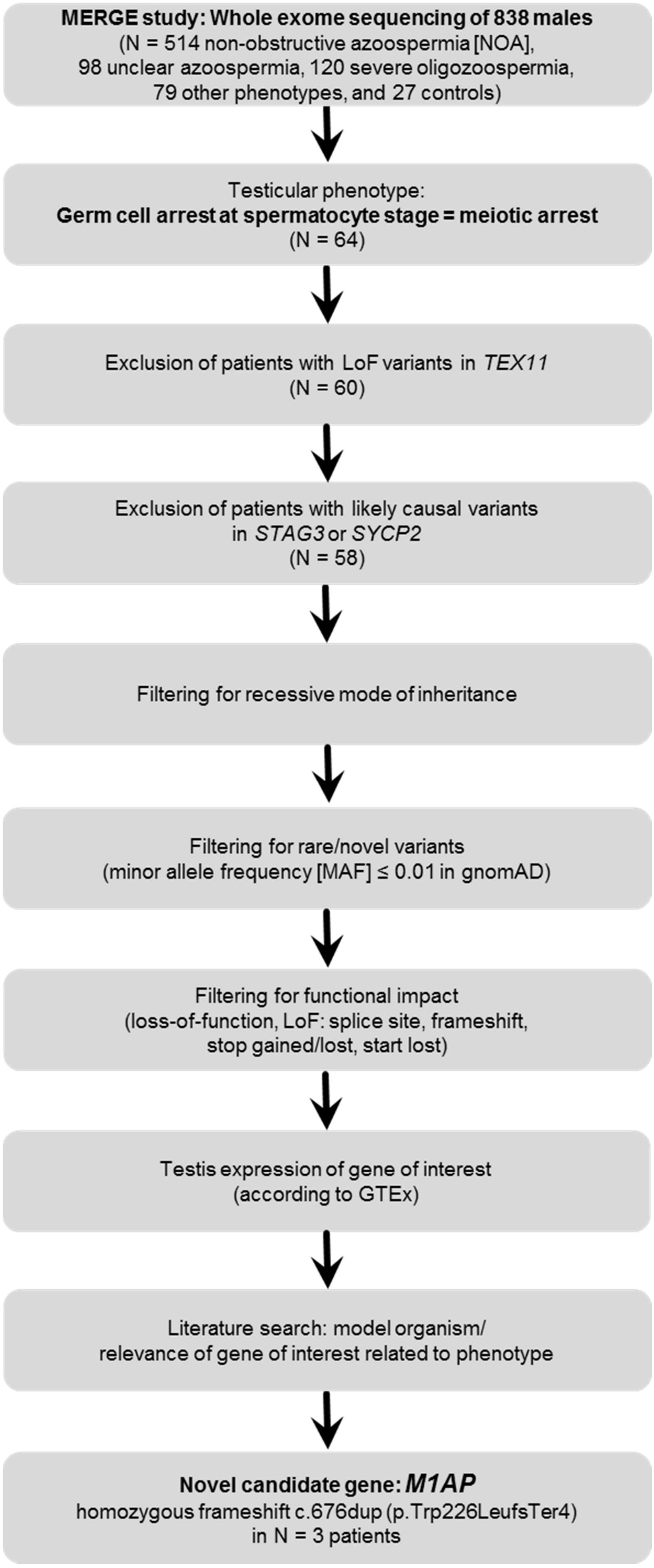

**Figure S2. Testis RNA analysis of patient M864.**

(**A**) PCR of testicular cDNA of exons 2-3 and 4-6 of a control and patient M864. Exons 2-3 and 4-6 were amplified. Expected band sizes were 146 bp (Exon 2-3) and 312 bp (Exon 4-6). + fertile control DNA, Mutation: patient M864 DNA, - RT: negative control without reverse transcriptase (RT), NTC: no template control. (**B**) Sanger sequencing of the PCR product amplified from cDNA of patient M864 was performed according standard procedures and validates the homozygous variant c.676dup in Exon 5.

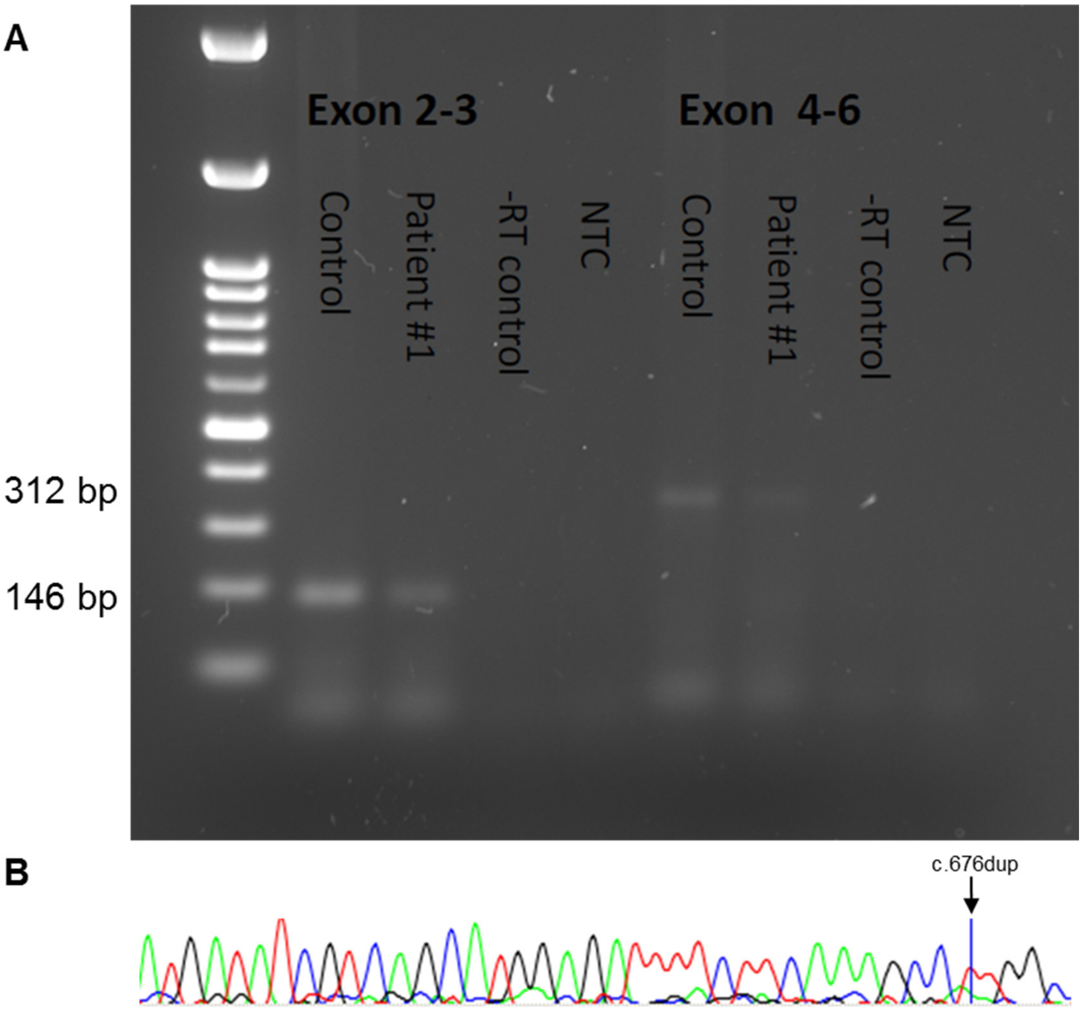

**Figure S3. Analysis of M1AP protein expression by immunohistochemical staining using #PA5-31627.**

(**A**) Low concentration of M1AP antibody (#PA5-31627, ThermoFisher Scientific, exemplarily depicted for 1:100) was evaluated and black arrow heads indicate spermatogonia, which potentially show weak but specific cytoplasmic protein expression. Additionally, unspecific antibody precipitation in control #2 can be observed (indicated by white arrow head). (**B**) Analysis of higher concentrations of M1AP antibody (exemplarily depicted for 1:50) lead to enhanced unspecific background signal and specific cytoplasmic protein expression primarily in spermatogonia (indicated by black arrow head). (**C**, **D**) MAGEA4 was used as a marker for early germ cells. (**E**, **F**) Omission of primary antibody (OC) is shown as representative technical control. Scale bars are indicated in each micrograph, respectively.

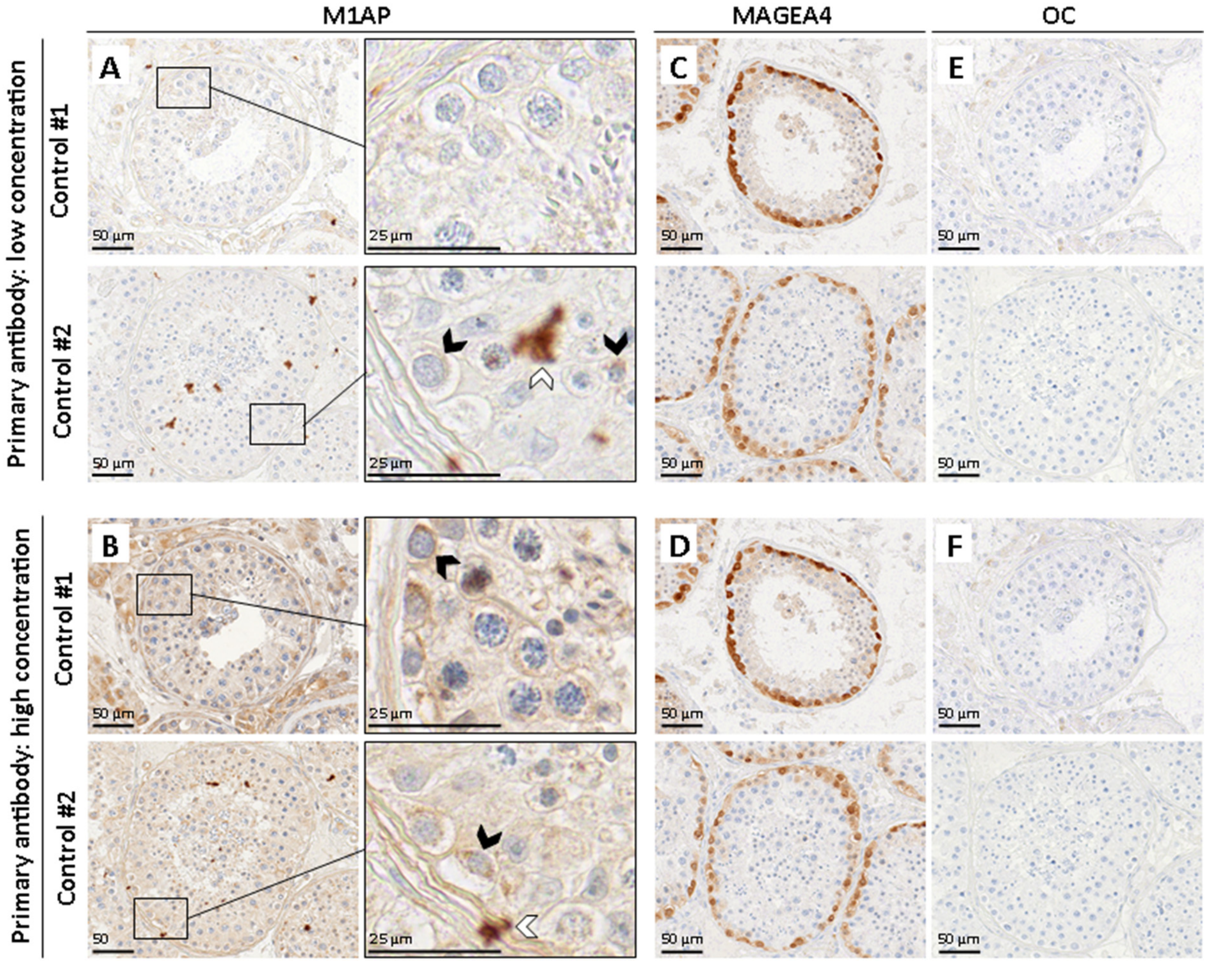

**Figure S4. Analysis of M1AP protein expression by immunohistochemical staining using #HPA045420.**

(**A**) Testis tissue from two patients with a homozygous duplication (c.676dup p.Trp226LeufsTer4) in *M1AP* resulting in bilateral meiotic arrest was stained with a commercially available M1AP antibody (#HPA045420, Sigma-Aldrich, exemplarily depicted for 1:100). Representative micrographs show cytoplasmic protein expression specifically in spermatogonia (indicated by black arrow head). (**B**) Sections from patients with full spermatogenesis were used as controls. The detected signal did not differ from *M1AP* mutated patients. (**C**, **D**) MAGEA4 was used as a marker for early germ cells. (**E**, **F**) Omission of primary antibody (OC) is shown as representative technical control. Scale bars are indicated in each micrograph, respectively.

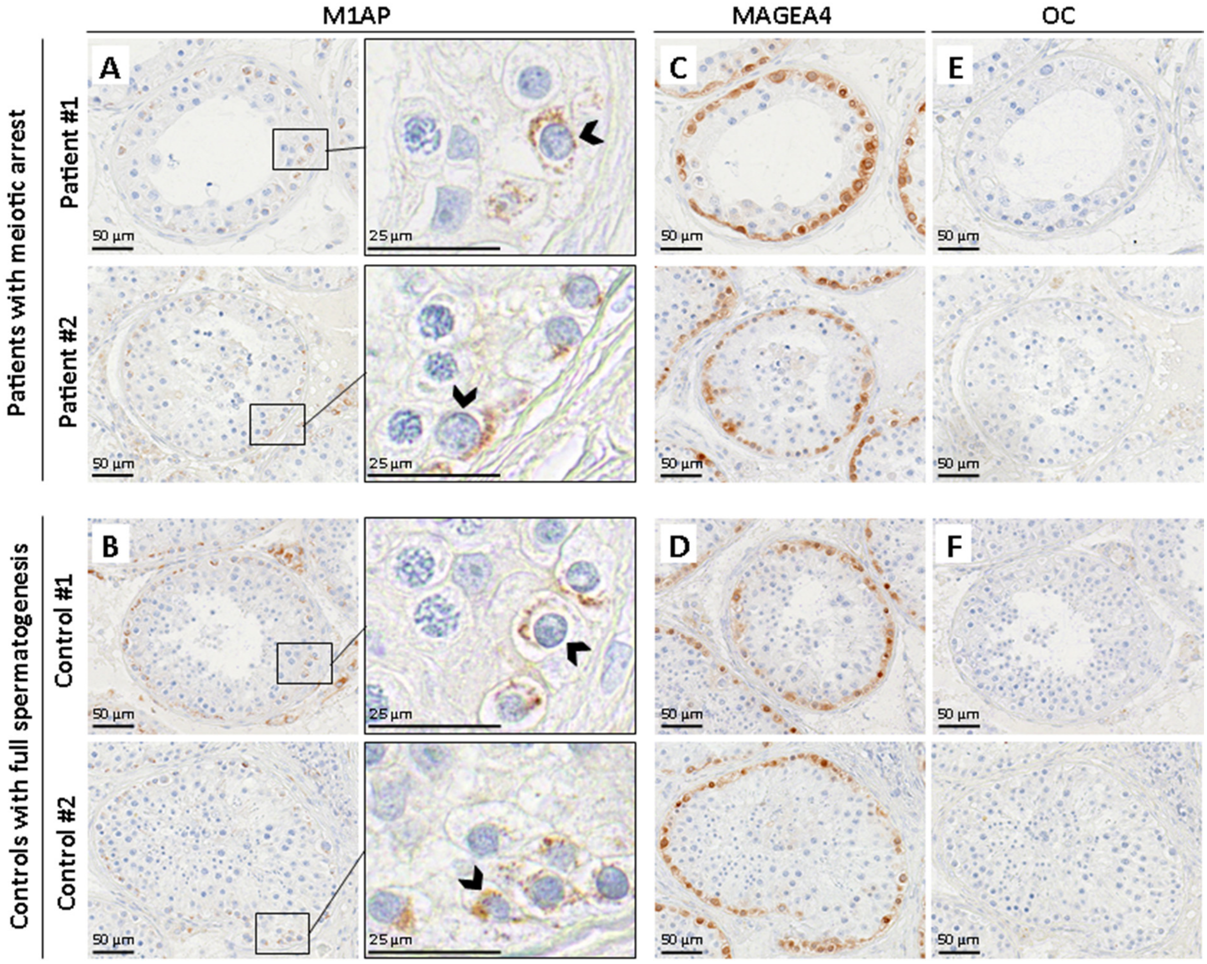

**Figure S5. Western blot analysis.**

Cytosolic (CL) and whole cell (WL) lysates from testis and kidney were obtained from fertile healthy donors. Western blots were stained with two different M1AP antibodies as indicated. Predicted molecular weight of M1AP is 59 kDa. The observed band size of almost 55 kDa is similar to those expected for PA5-31627 antibody by the manufacturer's instructions. In contrast, validation experiments of the Human Protein Atlas of HPA045420 antibody detect protein bands of ~30 kDa in different tissues and cell lines. Therefore, the specificity of both antibodies is questionable. Furthermore, M1AP is not expressed in kidney (GTEx, Human Protein Atlas).

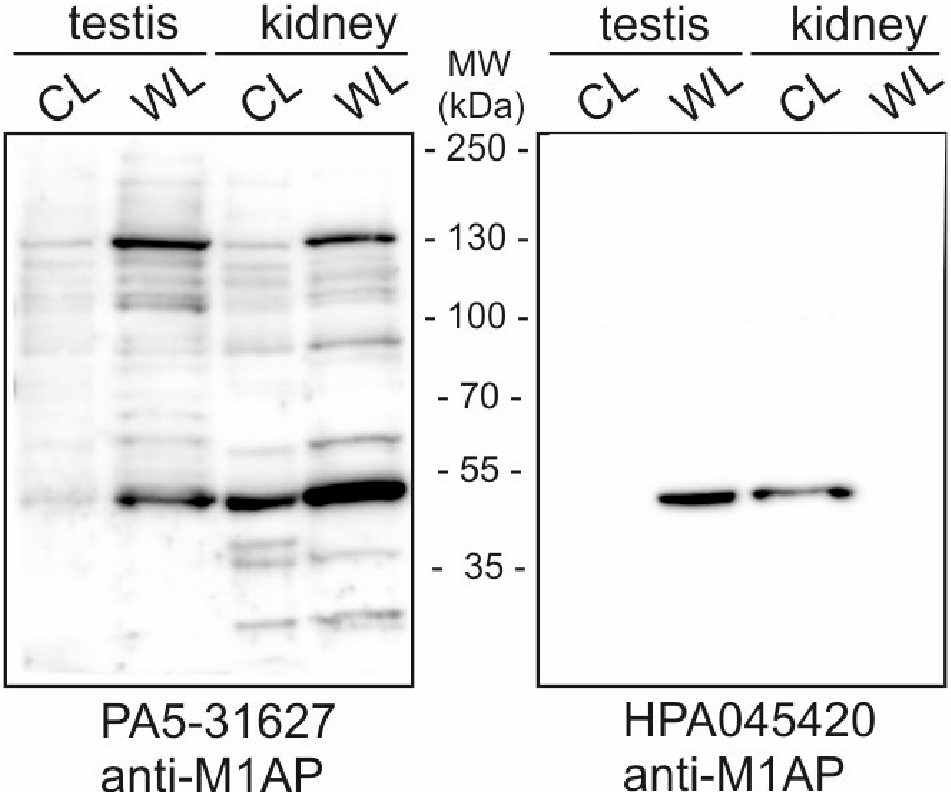

### Supplemental Methods

*Whole exome sequencing (WES) and bioinformatic analysis in patient RU01691*

Patient RU01691 came from a cohort of 99 patient-parent trios. These were cases of non-obstructive severe oligozoospermia (<5 million sperm/ml; N = 44) and azoospermia (no sperm in the ejaculate; N = 55). Exomes of the case-parent trios were enriched using the TruSeq Rapid Enrichment kit (Illumina, San Diego, CA) and the sequencing was performed on the NovaSeq 6000 Sequencing System (Illumina) at an average depth of 72x, with 91.7% of all targeted regions covered at least ten times.

*Whole exome sequencing (WES) and bioinformatic analysis in patient Y126 and P86*

Genetics of Male Infertility Initiative (GEMINI; https://gemini.conradlab.org/) is a multi-center consortium dedicated to identifying and describing the underlying genetic causes of male infertility. To date, N = 979 men with non-obstructive azoospermia have been studied using the whole-exome sequencing (WES) approach. In short, WES was performed at the McDonnell Genome Institute of Washington University (genome.wustl.edu) on Illumina HiSeq 4000 and using an in-house exome targeting reagent which captures 39.1 Mb of exome at an average coverage of 80x. The sequence reads were aligned to hg38 using bwa-mem, Picard and Genome Analysis Toolkit (GATK; https://software.broadinstitute.org/gatk) in an alternate contig-aware manner. Genotype calling was performed jointly for all samples using GATK tools and following their procedures of best practices. The resulting genotype callset was subjected to thorough quality control procedures, including but not limited to removing positions with high missingness rate (>15%) and removing samples with low coverage (<30x), high contamination (VerifyBamID freemix >5%) or low call rate (<85%). Individual genotypes with read depth (DP) <10x and genotype quality (GQ) <30 were further excluded from the data. All sequenced cases were screened for the known infertility causes such as Klinefelter syndrome, deleterious CFTR mutations, Y-chromosome microdeletions and large structural variation on sex chromosomes utilizing the WES genotype dataset.

In order to prioritize the deleterious lesions most likely disrupting the function of the respective genes and potentially leading to the disease phenotype, a modified version of the population sampling probability (PSAP) software (https://github.com/conradlab/PSAP) was applied to the WES genotype callset. The genomic coordinates of identified variation were lifted over to hg19 for the PSAP analysis. The list of prioritized variants was subsequently filtered by only including mutations with PSAP popScore values less than $10^{-4}$ and minor allele frequency <1% across all populations in the gnomAD database (v2.1.1, https://gnomad.broadinstitute.org/). As the study aims to identify rare DNA lesions most likely observed in few, if not singleton, cases, genes enriched for and positions found to be commonly affected by rare deleterious variation among the cases were excluded from the study.

*Whole exome sequencing (WES) and bioinformatic analysis in patient T1024*

Genomic DNA was extracted from peripheral venous blood using the QIAamp® DNA Mini Kit (QIAGEN, Ankara, Turkey). SureSelectXT Library Prep Kit was used for target enrichment. All procedures were carried out according to the manufacturer's protocols. Paired-end sequencing was performed on an Illumina NovaSeq system with a read length of 151. Base calling and image analysis were conducted using Illumina's Real-Time Analysis software. The BCL (base calls) binary is converted into FASTQ utilizing Illumina package bcl2fastq. All bioinformatics analysis performed on Sophia DDMTM platform which includes algorithms for alignment, calling SNPs and small indels (Pepper), calling copy number variations (Muskat) and functional annotation (Moka). Raw reads were aligned to the human reference genome (GRCh37/hg19). Variant filtering and interpretation performed on Sophia DDMTM. Integrative Genomics Viewer (IGV)16 was used to bam file visualization. In families with consanguineous marriages, the homozygosity mapping was carried out with HomSI.

*Attempt at 3D modelling of M1AP protein*

The longest coding transcript of *M1AP* (GENCODE: ENST00000290536.5, RefSeq: NM_001281296.1, NM_138804.3) translates to the protein sequence with UniProt identifier Q8TC57. This sequence was used to perform a basic local alignment search tool (BLAST) to the sequences of known protein structures in the protein data bank (PDB). Four structures were obtained that partially match the M1AP sequence.

1. PDB ID: 5CK3 (Chain B, D, and F) with sequence identity: 29%
2. PDB ID: 5CK4 (Chain A, and B) with sequence identity: 29%
3. PDB ID: 5CK5 (Chain A, B, C, and D) with sequence identity: 29%
4. PDB ID: 6EGC (Chain A) with sequence identity: 24%

The highest sequence identity of these was 29%. Using these template structures will create unreliable results for a homology model, as templates with below 30% identity are likely to lead to serious mispredictions.

*RNA analysis*

For RNA isolation, snap frozen testicular material of a patient carrying the *M1AP* variant (M864) and a control proband with full spermatogenesis was used. RNA isolation was conducted with the miR-Neasy Micro kit (217084, Qiagen, Hamburg, Germany) according to the manufacturer's protocol. 500 ng of RNA were used as starting material for cDNA synthesis employing the iScript cDNA Synthesis Kit (Biorad). Subsequently, 4 µl of cDNA were amplified via PCR reaction using the Qiagen Taq Polymerase Kit. Reactions were performed in 20 µl of total volume with primer concentrations of 20 pmol (sequences in Table S1) and 1 mM dNTPs. The PCR included an initial incubation step at 94°C for 2 min followed by 35 cycles at 94°C for 30 sec, 56°C for 45 sec and 72°C for 1 min as well as a final step at 72°C for 10 min. PCR products were evaluated using a 2% agarose gel.

*Attempt at M1AP immunohistochemical localization*

Testicular tissue sections of 3 µm thickness were prepared (SM2010R sliding microtome, Leica Biosystems, Nussloch, Germany) from two human control samples with complete spermatogenesis or patients with a homozygous duplication (c.676dup p.Trp226LeufsTer4) in *M1AP*, resulting in bilateral meiotic arrest. Sections were deparaffinized, rehydrated in a descending ethanol row, and rinsed with tap water. Heat-induced antigen retrieval was performed in citrate buffer (pH 6). After cooling to room temperature (RT), sections were washed with 1X Tris-buffered saline (TBS) prior to incubation with 3% $H_2O_2$ for 15 min at RT to inactivate endogenous peroxidases. Washing steps with distilled water and TBS followed. Nonspecific binding sites were blocked by applying 25% goat serum (#G6767-100ML, Sigma-Aldrich, Munich, Germany) in TBS containing 0.5% bovine serum albumin (BSA) for 30 min at RT in a humid chamber. Primary antibody incubation was performed overnight at 4°C in a humid chamber and different concentrations were evaluated for both M1AP antibodies (1:20, 1:50, 1:100, 1:200, 1:500, 1:1000 diluted in blocking solution, respectively). MAGE-A4, a marker for early human germ cells, was used as a positive control (kindly provided by Prof. G. C. Spagnoli, University Hospital of Basel, Switzerland). Respective IgG (#I5006, Sigma-Aldrich, Munich, Germany) and omission of primary antibody controls were included for each staining. On the following day, sections were washed again in TBS and incubated with corresponding secondary antibodies (goat anti-rabbit Biotin, #ab6012, Abcam, Cambridge, USA - 1:100, 1:200) for 1 h at RT in a humid chamber. After further washing with TBS, Streptavidin conjugated with horse-radish peroxidase (#S5512, Sigma-Aldrich, Munich, Germany) was diluted in blocking solution (1:500) and sections were incubated for 45 min at RT in a wet chamber. Subsequently, sections were washed with TBS and incubated with 3,3'-Diaminobenzidine tetrahydrochloride (DAB, #D3757, AppliChem, Darmstadt, Germany) for visualization of antibody binding. Staining was validated by microscopical acquisition and distilled water was used to stop the reaction. Counterstains were conducted using Mayer's hematoxylin (#109249, Merck Millipore, Darmstadt, Germany). Sections were rinsed with tap water, dehydrated, and mounted using Merckoglas® mounting medium (#103973, Merck Millipore, Darmstadt, Germany). Images were captured using the PreciPoint M8 Scanning Microscopy System (PreciPoint, Freising, Germany).

*Tissue preparation and Western blotting analysis*

Cytosolic lysates were prepared by mincing biopsy samples from human testis or kidney in IP buffer (1% Triton-X 100, 20 mM Tris-HCl [pH 7.5], 25 mM NaCl, 50 mM NaF, 15 mM Na4P2O7, and 1.5 mM EDTA) containing protease inhibitors (Complete; Roche, Mannheim, Germany). Samples were centrifuged at 10,000×g for 45 min at 4°C to pellet cell debris and nuclei. Cytosolic supernatants were removed and stored at −80°C until further use. For total cell lysates, tissue was minced in Laemmli buffer ((20% Glycerin, 125 mM Tris-HCl pH 6.8, 10% SDS, 0.2% Bromphenol blue, 5% β-Mercaptoethanol), boiled for 5 min at 95°C and then stored at -20°C. Sodium dodecyl sulfate polyacrylamid gel electrophoresis (SDS-PAGE) and semidry Western blotting analysis was performed using standard techniques. Primary anti-M1AP antibodies were from ThermoFisher (PA5-31627) and Sigma (HPA045420), respectively. Signals were detected using secondary peroxidase-coupled antibodies and enhanced chemiluminescence (ECL).
